# Rediscovery and redescription of the endangered *Hypostomus subcarinatus* Castelnau, 1855 (Siluriformes: Loricariidae) from the rio São Francisco basin in Brazil, with comments on the urban water conservation

**DOI:** 10.1101/458604

**Authors:** Cláudio Henrique Zawadzki, Iago de Souza Penido, Tiago Casarim Pessali

## Abstract

*Hypostomus subcarinatus* Castelnau, 1855 is rediscovered in the Pampulha lake, an urban lake pertaining to the rio das Velhas basin in the rio São Francisco system in the state of Minas Gerais, southeastern Brazil. Herein, *H. subcarinatus* is redescribed and its diagnosis from the congeners is established due to characters such as blue tan dorsal fin in live specimens, slender bicuspid teeth, dentaries angled more than 90 degrees, moderate keels along lateral series of plates, small roundish dark spots, one plate bordering supraoccipital, by having nuptial odontodes mainly on pectoral, dorsal and caudal-fin rays, and longer anal-fin unbranched ray. The rediscovery of *H. subcarinatus* after more than 160 years after its original description was one unexpected event, because the Pampulha lake is an artificial, shallow and polluted urban lake. The lake is located in the downtown of municipality of Belo Horizonte, the third largest urban agglomeration in Brazil with a population exceeding 5.9 million inhabitants. In the light of this finding we address the importance of urban body waters to maintenance of fish biodiversity in the neotropics.

## Introduction

The loricariid *Hypostomus subcarinatus* was described by Castelnau [1] from a vague type locality stated as “des rivière de la province des Mines” [streams from the state of Minas Gerais]. Therefore, it was hypothetically asserted to the Eastern Brazilian coastal drainage and to the rio São Francisco basin [2]. However, despite some ichthyological survey efforts in these systems [3,4], none scientifically record of *H. subcarinatus* was undoubtedly stated for more than 160 years. This historical *H. subcarinatus* lack of records lead to some hypothesis, a) an erroneous locality designation in the original description by Castelnau; b) species rarity or endemicity to specific locations; c) several ongoing populational extinction processes; or d) imprecise identifications.

In 2014 it was accomplished a fish environmental monitoring of the Pampulha lake, an artificial shallow and polluted urban lake pertaining to the rio São Francisco basin system and located in the downtown of municipality of Belo Horizonte, Minas Gerais State, southeastern Brazil. Unexpectedly, in the Pampulha lake seven large specimens of the catfish *Hypostomus* were captured. Subsequent specimens examination did not allow to recognize them to any of the commonly found species of *Hypostomus* from the rio São Francisco basin. However, in comparison to *Hypostomus* original descriptions, as well as to types series of *Hypostomus* from worldwide scientific fish museums we finally recognized the specimens as the Castelnau’s (1855) “lost” *Hypostomus subcarinatus.* In this work we redescribe the species and discuss about the importance of the conservation of urban water body.

## Material and Methods

Fishes were collected under permits from the Instituto Chico Mendes de Conservação da Biodiversidade – ICMBio n. 9101-1/2017. Captured individuals were anaesthetized and sacrificed by immersion in eugenol (active ingredient: phenolic eugenol, 4-Allyl-2-methoxyphenol-C10H1202, derived from stems, flowers and leaves of *Eugenia caryophyllata* and *Eugenia aromatica* trees) [5], fixed in 10% formalin solution and later preserved in 70% ethanol. These procedures are in accordance to the ‘Ethical Principles in Animal Research’ guidelines adopted by the Brazilian College of Animal Experimentation (COBEA). Measurements and counts of bilaterally symmetrical features were taken from the left side of the body, whenever possible. Measurements were taken using a digital caliper to the nearest 0.1 mm. Methodology and terminology of measurements follows Boeseman [6], modified by Weber [7] and Zawadzki *et al.* [8]. Plate counts and bone nomenclature follow Schaefer [9], modified by Oyakawa *et al.* [10]. Standard length (SL) is expressed in millimeters and all other measurements are expressed as percents of standard length or head length (HL), unless otherwise noted. Institutional abbreviations of material deposited follow Fricke & Eschmeyer [11]. The species conservation status was calculated through the criteria by the International Union for Conservation of Nature (IUCN standards and petitions subcommittees, 2017 [12]) guideline. The Extent of Occurrence (EOO) was calculated by the minimum convex polygon method, using the software Google Earth Pro.

## Results

*Hypostomus subcarinatus*, Castelnau, 1855 (Figs. 1, 2 and 3, Table 1)

**Figure 1.**
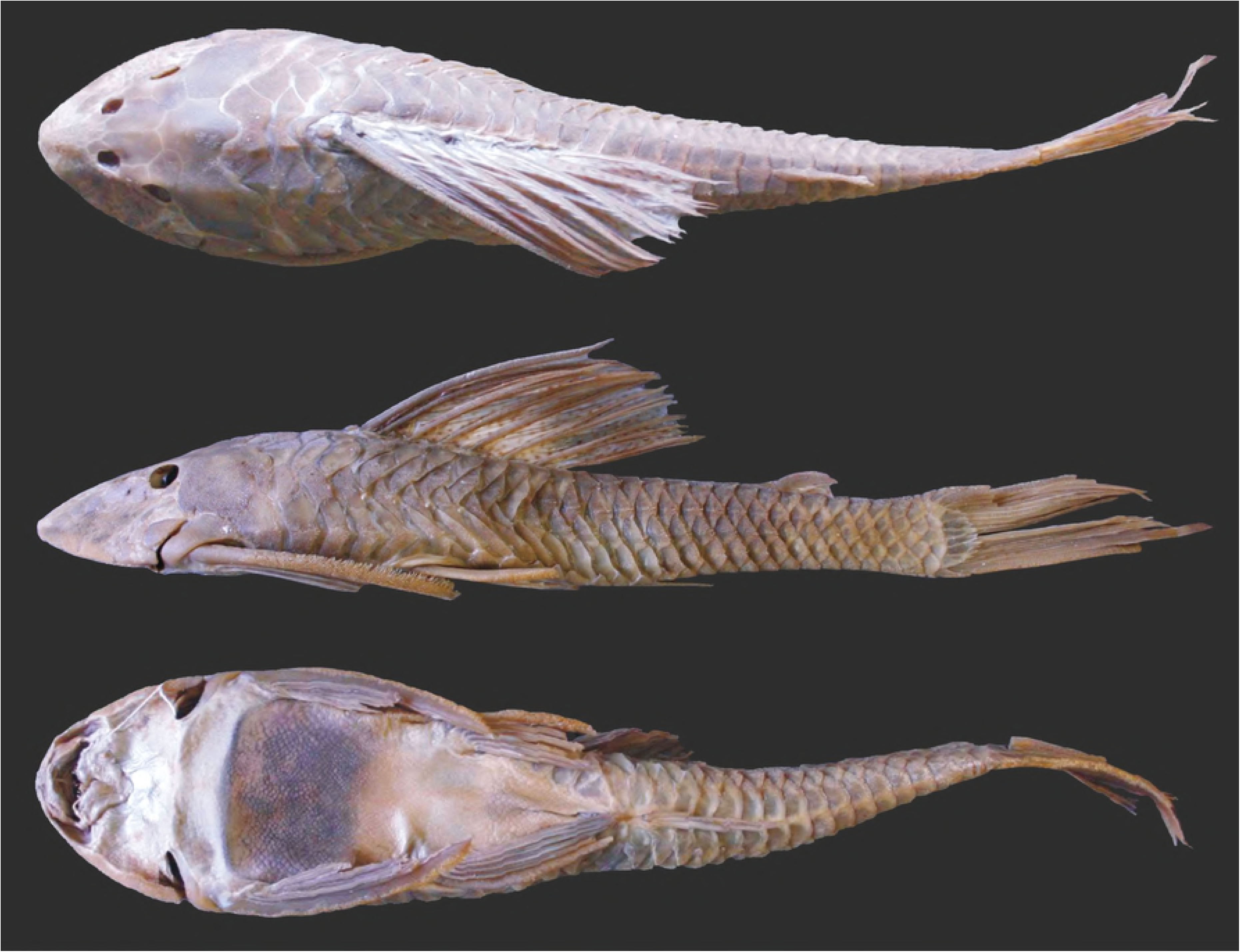
*Hypostomus* subcarinaus, MNHN A, 9575, 241.8 mm SL, holotype, Brazil, Province de Mines [estado de Minas Gerais].

**Figure 2.**
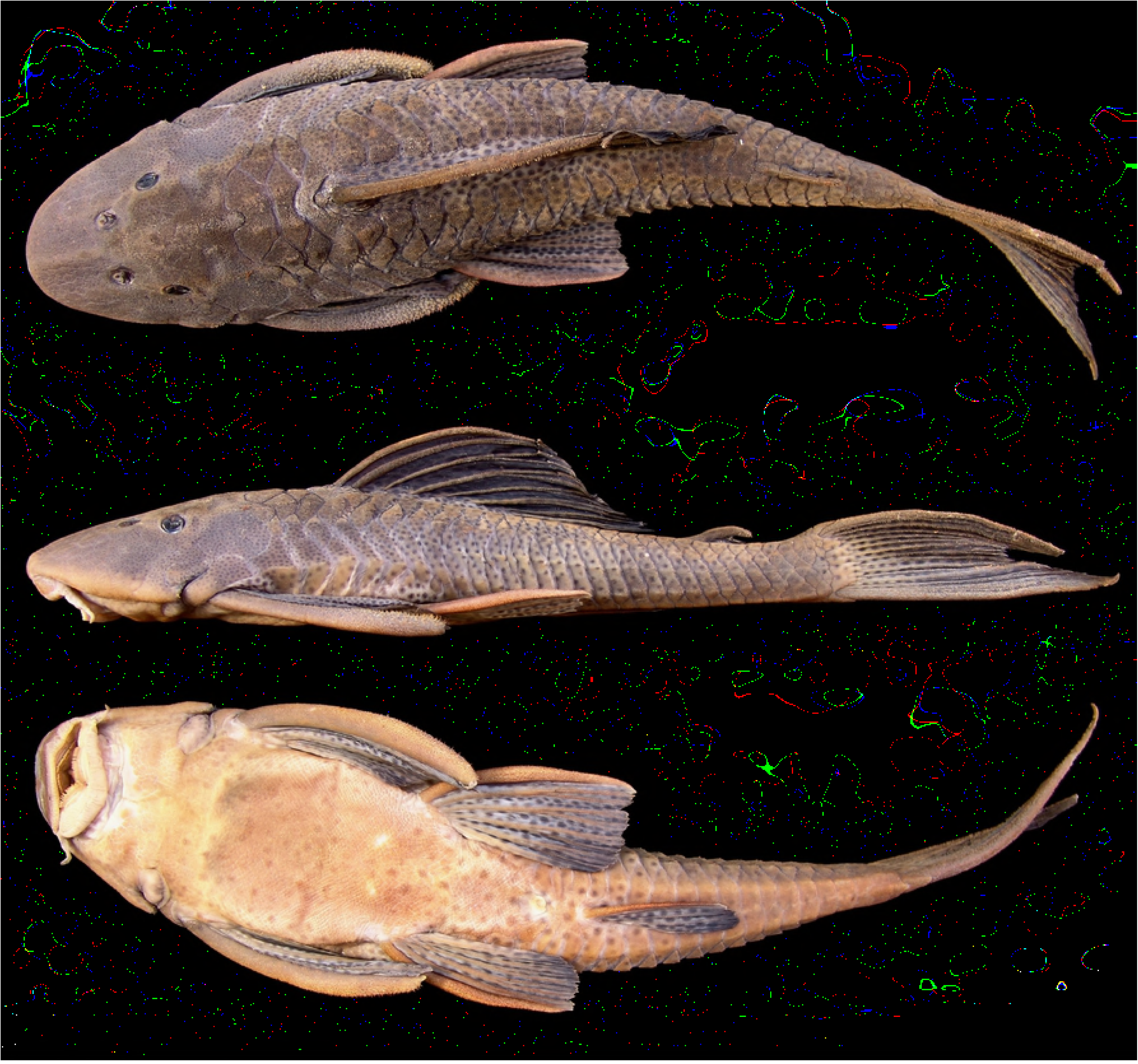
*Hypostomus subcarinatus* MCNIP 1103, 249.5 mm SL. Pampulha lake, Belo Horizonte, Minas Gerais State, Brazil.

**Figure 3.**
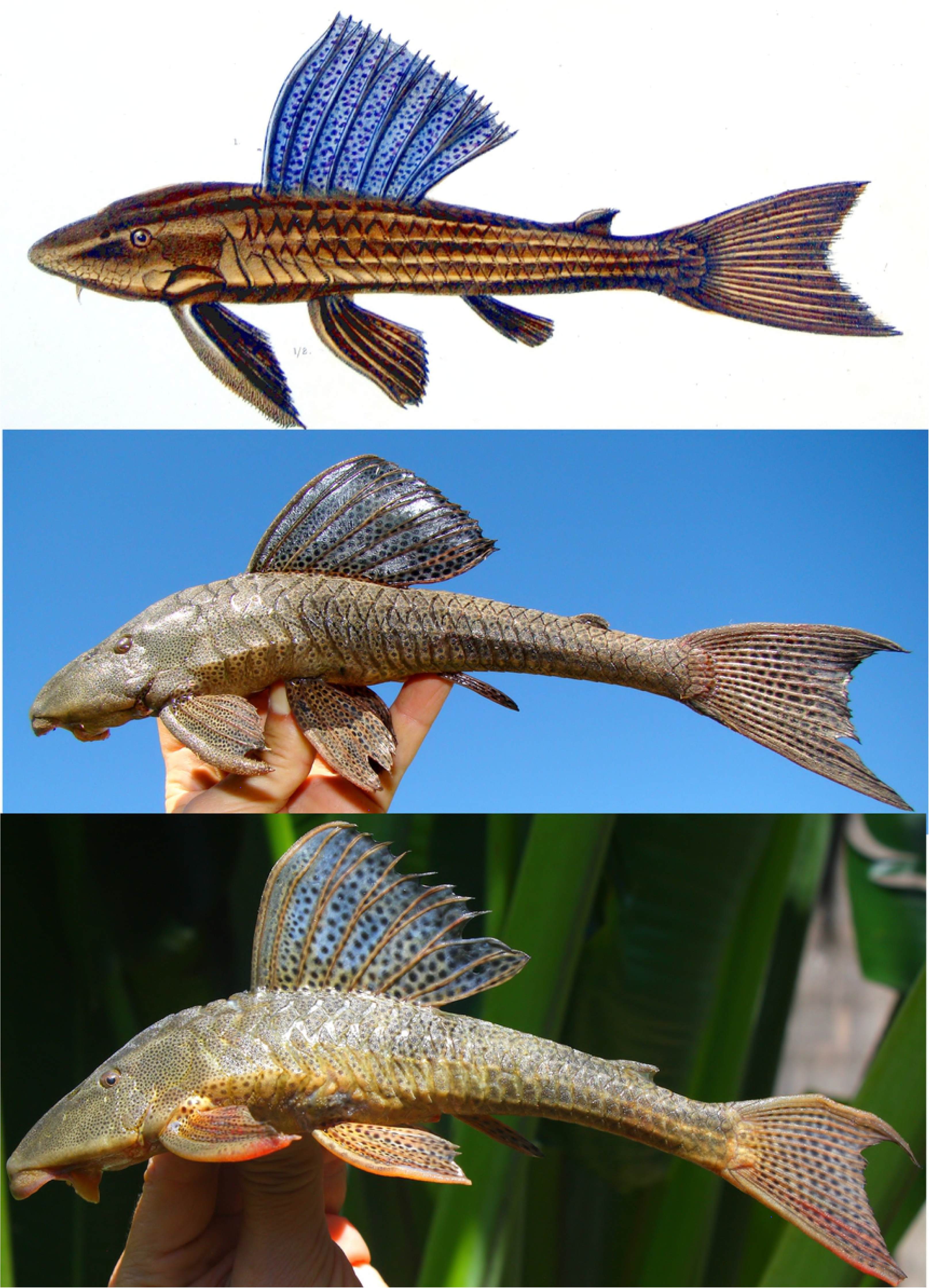
The Castelnau’s (1855) drawing of *Hypostomus subcarinatus*, MNHN A 9575, 241.8 mm SL, holotype, Brazil, Province de Mines [state of Minas Gerais], is depicted in the upper picture and compared to two live specimens photographed immediately after capture: MCNIP 1761, middle picture 227.2 mm SL and lower picture 196.5 mm SL, both from the Pampulha lake, Belo Horizonte, Minas Gerais State, Brazil.

**Table 1.** Morphometrics and counts of *Hypostomus subcarinatus*. N = 24 specimens (range not including holotype). SD = Standard deviation.

|  | holotype | range | mean | SD |
| --- | --- | --- | --- | --- |
| Standard length (mm) | 241.08 | 158.1–308.9 | 215.5 | 38.4 |
| **Percents of SL** |  |  |  |  |
| Predorsal length | 35.7 | 33.8–37 | 35.4 | 1.0 |
| Head length | 29.1 | 26.9–29.7 | 28.2 | 0.8 |
| Cleithral width | 25.6 | 23.4–26.6 | 24.8 | 0.8 |
| Head depth | 15.8 | 15.4–17.8 | 17.1 | 0.5 |
| Interdorsal distance | 23.7 | 20.3–23.6 | 22.3 | 0.9 |
| Caudal peduncle length | 34.9 | 31.8–36 | 34.4 | 1.1 |
| Caudal peduncle depth | 8.3 | 7.1–8.4 | 7.7 | 0.3 |
| Dorsal-spine length | 30.4 | 22.7–33.1 | 27.8 | 2.7 |
| Thoracic length | 23 | 20.7–24.4 | 22.8 | 1.0 |
| **Percents of head length** |  |  |  |  |
| Cleithral width | 87.9 | 84.6–92.7 | 87.7 | 2.4 |
| Head depth | 54.4 | 57.5–63.2 | 60.5 | 1.6 |
| Snout length | 60.3 | 57.4–62.4 | 59.8 | 1.4 |
| Orbital diameter | 11.8 | 11.1–13.7 | 12.5 | 0.7 |
| Interorbital width | 38.5 | 33.1–39.4 | 37.1 | 1.6 |
| Mandibular width | 15.7 | 12.4–14.7 | 13.8 | 0.7 |
| **Other percents** |  |  |  |  |
| Orbital diameter in snout length | 19.6 | 17.8–23.9 | 20.9 | 1.6 |
| Orbital diameter in interorbital length | 30.7 | 29.5–38.4 | 22.7 | 2.9 |
| Mandibular length in interorbital length | 40.7 | 31.4–43.8 | 37.2 | 2.6 |
| Dorsal-spine length in predorsal length | 85.1 | 66.3–93.5 | 79.1 | 7.3 |
| First pectoral-fin ray length in predorsal length | 75.8 | 70.4–79.5 | 73.8 | 2.8 |
| Ventral caudal-fin ray length in predorsal length | 88.4 | 91.6–108.3 | 97.4 | 4.6 |
| Adipose-fin length in caudal peduncle depth | 85.5 | 72.–106.6 | 87.9 | 10.8 |
| Caudal peduncle depth in caudal peduncle length | 23.8 | 20.4–25 | 22.5 | 1.4 |
| Mandibular width in cleithral width | 17.8 | 13.3–17.1 | 15.7 | 0.9 |
| Interdorsal length in dorsal-fin base | 98.3 | 82.6–99.1 | 91.9 | 4.2 |
| Lower lip length in lower lip width | 33.1 | 22.2–34.5 | 27 | 3.3 |
| **Counts** |  |  |  |  |
| Median plates series | 28 | 28–32 | 30 |  |
| Plates bordering supraoccipital | 1 | 1–1 | 1 |  |
| Predorsal plates | 3 | 3–3 | 3 |  |
| Dorsal plates below dorsal fin bases | 8 | 8–8 | 8 |  |
| Plates between dorsal and adipose fins | 9 | 7–9 | 8 |  |
| Plates between adipose and caudal fins | 6 | 6–7 | 6 |  |
| Plates between anal and caudal fins | 14 | 14–16 | 15 |  |
| Premaxillary teeth | 34 | 37–53 | 43 |  |
| Dentary teeth | 34 | 36–54 | 44 |  |

### Type-specimens

Holotype. MNHN A. 9575, 241.8 mm SL, des rivière de la province des Mines [streams from the state of Minas Gerais].

### Material analized

All from Brazil, Minas Gerais State: rio São Francisco basin: MCNIP 1103, 7, 158.1-249 mm SL, municipality of Belo Horizonte, Pampulha lake, tributary of córrego da Onça, rio das Velhas basin, 19°50’30”S 43°59’38”W, 30 Jan 2014, A. A. Weber & D. Gontijo. NUP 20229, 7, 164.9-248.9 mm SL, municipality of Belo Horizonte, Pampulha lake, tributary of córrego da Onça, rio das Velhas basin, 19°50’30”S 43°59’38”W, 23 Dec 2017, I. S. Penido & T. C. Pessali. MCNIP 1761, 7, 191.1-308.8 mm SL, municipality of Belo Horizonte, Pampulha Lake, tributary of córrego da Onça, rio das Velhas basin, 19050’30”S 43059’38”W, 15 Apr 2016, I. S. Penido, C.H. Zawadzki, F. M. Azevedo & T. C. Pessali.

### Diagnosis

*Hypostomus subcarinatus* is distinguished from all congeners by having blue tan dorsal fin in living specimens (vs. not having blue tan dorsal fin). Additionally, *H. subcarinatus* is diagnosed from the species of the *H*. *cochliodon* group by having slender viliform bicuspid teeth (vs. robust spoon-shaped teeth) and by having dentaries angled to each other more than 90 degrees (vs. dentaries angled from 80 to 90 degrees); from the remaining congeners except *H. affinis*, *H. ancistroides*, *H. argus*, *H. aspilogaster*, *H. borellii*, *H. boulengeri*, *H. carinatus*, *H. careopinnatus*, *H. commersoni*, *H. corantijni*, *H. crassicauda*, H. *delimai*, H. *dlouhyi, H. faveolus, H. formosae, H. gymnorhynchus*, H. *hemiurus*, H. *hoplonites, H. interruptus, H. micromaculatus*, *H. niceforoi, H. nigrolineatus*, *H. pantherinus, H. paucimaculatus*, *H. piratatu*, *H. plecostomus*, *H. punctatus*, *H. pusarum*, *H. rhantos*, *H. scabriceps*, *H. seminudus*, *H. tapijara*, *H. velhochico* and *H. watwata*, by having moderate keels along the five lateral series of plates (vs. lacking keels); from *H. affinis*, *H. ancistroides, H. argos, H. aspilogaster, H. borellii, H. boulengeri*, *H. carinatus, H. careopinnatus, H. commersoni, H. corantijni, H. crassicauda*, *H. delimai, H. dlouhyi, H. faveolus, H. formosae, H. gymnorhynchus, H. hemiurus, H. hoplonites, H. interruptus, H. micromaculatus, H. niceforoi, H. nigrolineatus, H. pantherinus, H. paucimaculatus, H. piratatu, H. plecostomus*, *H. punctatus*, *H. pusarum*, *H. rhantos*, *H. scabriceps*, *H. seminudus*, *H. tapijara*, *H. velhochico* and *H. watwata* by having more elongate and slender body, having a longer anal-fin unbranched ray, anal-fin unbranched ray length almost or equal to nostril length, that is, the distance from anterior margin of snout to anterior edge of eye (vs. shorter anal-fin unbranched ray, its length equal to nare length, that is, the distance from the anterior margin of nostril to nare).

### Description

Morphometric data in Table 1. Overall view of body in Figs. 1, 2 and 3. Head moderately depressed and slightly compressed. Snout and anterior profile of head slightly pointed in dorsal view. Eye of small size, dorsolaterally positioned. Dorsal margin of orbit not raised. Greatest body width at cleithrum, narrowing from dorsal-fin region to caudal-fin origin. Dorsal profile of head convex from snout tip to vertical through interorbital region, forming angle of about 40° with ventral region of head; slightly convex from that point to dorsal-fin origin; straight from that point to caudal peduncle end; rising to procurrent rays of dorsal fin. Ventral profile almost straight from snout tip to insertion of pelvic-fin unbranched ray; tapering slightly straight from pelvic-fin insertion to first ventral caudal-fin procurrent ray. Anterior portion of caudal peduncle rounded with its dorsal surface compressed; posterior portion ellipsoid. Mesethmoid forming weak longitudinal bulge from snout tip to nares. Supraoccipital bone with slightly-developed median ridge and short posterior process bordered by single plate. Weak bulge originating lateral to nares, passing through supraorbital, and extending as ridge along dorsal portion of pterotic-supracleithrum. Opercle large, its horizontal length equal to distance between nares, with thin skin layer surrounding its ventral edges to subocular cheek plates. Oral disk round, moderate in size; its margins smooth. Lower lip far from reach transverse line through gill openings; ventral surface with two to three transverse dermal flaps posteriorly margining each dentary rami; short naked area followed by larger area with numerous small papillae decreasing in size distally. Maxillary barbel moderately long, slightly larger than eye to nare distance; mostly free from lower lip. Odontodes present on anterior surface of upper lip, just below snout. Dentaries moderate to strongly angled, averaging from 90° a 100° between left and right dentary rami. Teeth viliform, bicuspid with lateral cusp smaller than mesial cusp; crowns bent ventrally. Internally to mouth, transversal areas of short papillae bordering each premaxillary and dentary teeth rami. Median buccal papilla present and well developed.

Body covered with five rows of dermal plates with moderately-developed odontodes, except on base of dorsal fin and small naked area on snout tip. Predorsal region with very slight median keel. Dorsal, mid-dorsal, mid-ventral, and ventral series of plates with moderate keels. Median series with weakly developed keels; bearing continuous lateral line. Ventral series bent ventrally. Ventral surface of head covered with platelets, except for region beneath lower lip. Abdomen covered with minute platelets in specimens larger than 90 mm SL, with exception of very small areas around pectoral- and pelvic-fin insertions. Distal portion of pterygiophore exposed.

Dorsal fin II,7, its origin at vertical just posterior midpoint between pectoral- and pelvic-fin insertions; first spine present as V-shaped spinelet. Distal margin of dorsal fin slightly convex; tip of last dorsal-fin ray from two to three plates to reach adipose-fin spine. Adipose-fin spine compressed and slightly curved inward. Pectoral fin I,6, its distal border straight. Pectoral-fin spine slightly curved inward, covered with moderately developed odontodes. Odontodes curved inward, more developed along distal portions of spine, particularly in larger specimens; emerging from swollen papillae. Tip of adpressed pectoral fin reaching to basal one-fourth to one-fifth of adpressed pelvic-fin unbranched ray. Pelvic fin i,5, its distal border straight to slightly convex; its adpressed unbranched ray surpassing one to two plates anal-fin origin. Anal fin i,4, its tip reaching to seventh plate after its origin; its distal margin straight. Caudal fin i,14,i, its margin falcate, with ventral lobe longer than dorsal.

### Color in alcohol

Overall ground color of dorsal and ventral regions of body and fins grayish-brown (Figs. 1 and 3). Head, trunk and fins covered by numerous small dark brown spots except on lower lip. Spots very small, numerous, close together and inconspicuous in head; increasing in length towards posterior region of body; spots more conspicuous on fins and dorsolateral regions of trunk. Spots on ventrolateral regions of trunk usually inconspicuous. Ventral surface of body usually with faded dark spots; conspicuousness variable among specimens. All fins with many small dark spots; spots irregularly distributed on spines and either on unbranched and branched rays. Some specimens with five faded oblique dark bars on dorsum, first bar on posterior portion of head, stronger at middle of orbit, second bar at first dorsal-fin branched rays, third bar at last dorsal-fin branched ray, fourth bar at anterior region of adipose fin and fifth bar at procurrent caudal-fin rays. Ventral surface of body slightly clearer than dorsal surface.

### Color in life

Color pattern of living specimens similar to preserved ones, except for more green-brownish background, black and more conspicuous spots and dorsal fin with blue tan (Fig. 2).

### Sexual dimorphism

No sexual dimorphism was observed among the specimens.

### Distribution

*Hypostomus subcarinatus* is known from one locality (Figs. 4 and 5); the Pampulha lake, an eutrophic reservoir, in the rio das Velhas basin, city of Belo Horizonte. Apparently the distribution of *H. subcarinatus* are restricted in this locality. However, more efforts of collections are needed.

**Figure 4.**
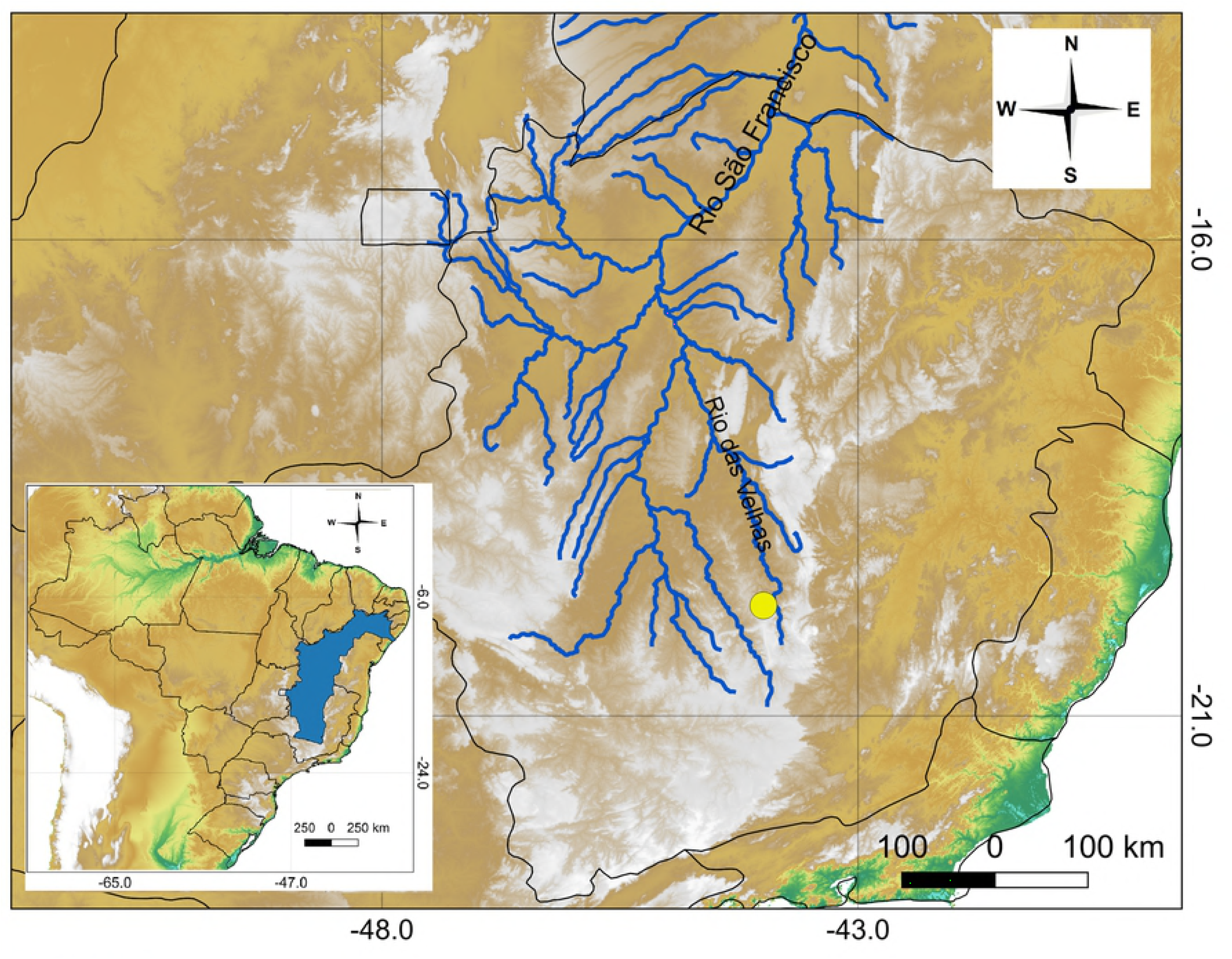
Geographical distribution of *Hypostomus subcarinatus;* (yellow circle = Pampulha lake). Blue shaded area and lines means the rio São Francisco basin.

**Figure 5.**
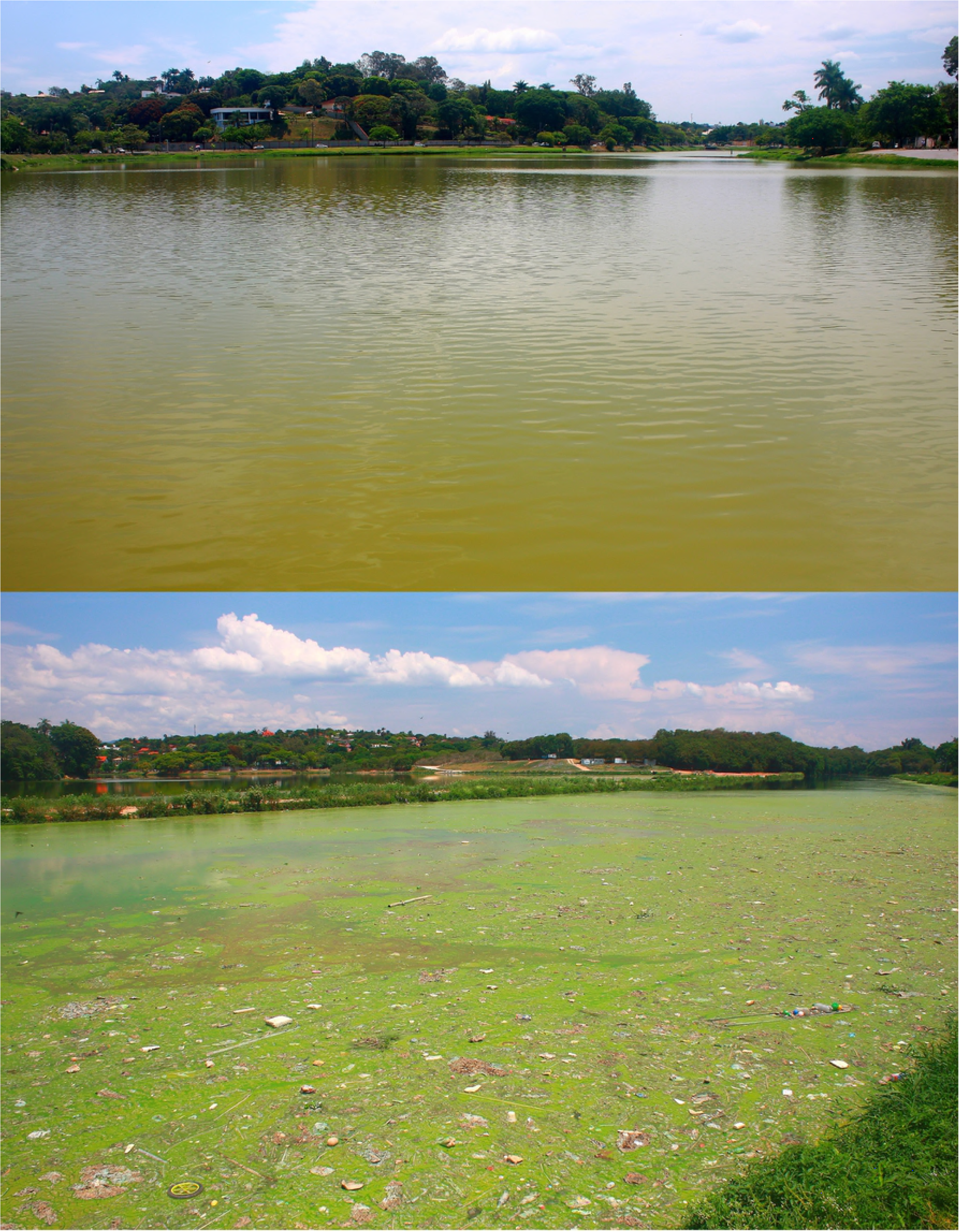
The Pampulha lake, at downtown of the city of Belo Horizonte, Minas Gerais State, Brazil. The habitat of *Hypostomus subcarinatus*.

### Habitat and conservation status

*Hypostomus subcarinatus* was up to now only found in the Pampulha lake, a silted and polluted urban reservoir (Fig. 5). The Pampulha lake was formed in 1938 to water supply to the city of Belo Horizonte. Since 1970 the reservoir has became quite eutrophic due to the receipt of domestic and industrial effluents from the city, causing recurrent cyanobacteria blooms [13]. Friese *et al.* [14] found significant values of heavy metals in lake sediments. As *Hypostomus* are known to be detritivorous fishes they probably assimilate considerable proportions of those metals as Veado *et al.* [15] found in the onivorous ciclhid *Oreochromis niloticus* in the Pampulha lake. Despite some ichthyologic survey efforts *H. subcarinatus* were up to now not collected in the surroundings of the lake. Therefore, *H. subcarinatus* with an estimated occupancy area of 1.96 km^2^ is herein considered critically endangered (CR) according to IUCN criterion, becoming the first threatened species of the genus. *Hypostomus subcarinatus* occurs syntopically to three alien cichlids in the lake, *Coptodon rendalli*, *Oreochromis niloticus, Parachromis managuensis.*

## Discussion

Concerning external morphology, the most similar species to *Hypostomus subcarinatus* are the eastern Brazilian drainage dwellers *H. affinis*, *H. interruptus*, *H. nigrolineatus*, *H. punctatus* and *H. scabriceps*. All species has elongate and narrow body with small to medium-sized dark spots and weak to moderate keels along lateral series of plates. Besides the dorsal-fin blue tan, *H. subcarinatus* is distinguished from these congeners due to be even more elongate and lower, having a longer anal-fin unbranched ray. Anal-fin unbranched ray length is almost or equal to nostril length vs. equal to the nostril-nares distance. Additionally, *H. subcarinatus* is also distinguished from *H. nigrolineatus* by having unorganized dark spots versus dark spots horizontally aligned to form conspicuous stripes on laterals of trunk.

Few papers dealt with *Hypostomus* from the rio São Francisco basin and its taxonomic issue is far from being well known [16]. Most *Hypostomus* records for the basin are data from dam construction monitoring programs, not resulting in ecological or taxonomic citations on scientific papers. However, several specimens of *Hypostomus* are deposited in ichthyological collections, mainly in the Museu de Ciências Naturais da PUC-MG, ICT-UFMG, Museu Nacional and at the Museu de Zoologia, Universidade de São Paulo, among others. Except the specimens from the Pampulha lake, *H. subcarinatus* were not recognized from *Hypostomus* samples at these collections.

Finding the native *H. subcarinatus* in the Pampulha lake at downtown of the city of Belo Horizonte, the third largest metropolis of Brazil with more than 5.9 millions inhabitants, was indeed a quite unexpected event. This is a fish larger than 300 mm in total length rediscovered more than 160 years after its original description and last citation. The individuals were found in the shallow, polluted urban lake, which is a significant ecological event. Some papers dealt with ecological surveys in urban neotropical streams [17, 18], but focusing fish conservation on urban neotropical artificial lake is an underestimated issue. Our findings highlight the importance that taxonomic focused scientific surveys in such a highly vulnerable water bodies can reveal important data to vertebrate conservation purposes. Urban lakes are frequently dragged, canalized, dried, and cleaned, to a series of reasons for human purposes. Our finding shows that in Netropical systems, even bad smelling urban waters as the Pampulha lake can harbor rare and endangered large fish, deserving conservation management.

### Comparative material

All from Brazil, unless noticed: *Hypostomus alatus:* Minas Gerais State, rio São Francisco basin: NUP 9119, 1, 110.1 mm SL, rio Curimataí. NUP 9829, 5, 139.0–177.4 mm SL, rio das Velhas. NUP 9837, 4, 124.4–217.6 mm SL, rio Cipó.

#### Hypostomus ancistroides

São Paulo State, rioTietê basin. LBP 2520, 2, 111.4–112.2 mm SL, rio Tietê. MCP 28309, 1, 138.0 mm SL, rio Piracicaba. MCP 28310, 3, 111.0–149.0 mm SL, rio Piracicaba.

MZUSP 2131, 4, 95.6–165.1 mm SL, rio Tatuí. NUP 64, 2, 55.0–74.6 mm SL, rio Capivara. NUP 4012, 3, 75.1–86.1 mm SL, rio Ipanema. NUP 4016, 5, 89.1–133.6 mm SL, rio Corumbataí.

#### Hypostomus aspilogaster

Rio Grande do Sul State, rio Uruguai basin. ANSP 21781, 1, 204.0 mm SL, lectotype (designated by Reis *et al.*, 1990), rio Jacuí. ANSP 21782, 3, 210.6–190.0 mm SL, paralectotypes, rio Jacuí. NUP 4355, 1, 155.0 mm SL, rio Ibicuí da Armada.

#### Hypostomus borellii

Bolivia. Rio Paraguai basin. BMNH 1897.1.27.19, 1, 153.1 mm SL, syntype, río Pilcomayo.

#### Hypostomus boulengeri

Mato Grosso State, rio Paraguai basin. NUP 414, 3, 165.8–175.6 mm SL; NUP 3273, 8, 110.0–166.0 mm SL; NUP 8695, 1, 170.0 mm SL, rio Manso. NUP 1078, 2, 210.0–220.0 mm SL, rio Manso Reservoir. NUP 8692, 1, 190.0 mm SL, rio Quilombo, rio Manso basin.

#### Hypostomus brevicauda

Bahia State. BMNH 1864.1.19.16–17, 2, 189.0–196.1 mm SL, syntypes. MCP 36709, 3, 52,7–125.4 mm SL, córrego Traíra, municipality of Camacã. MZUSP 111259, 4, 40.5–113.4 mm SL, rio Gongogi, tributary of rio de Contas.

#### Hypostomus carinatus

Amazonas State, Rio Amazonas basin INPA 1198, 2, 176.7 mm SL, rio Trombetas. INPA 2535, 1, 182.6 mm SL and INPA 2541, 1, 191.9 mm SL, Rio Uatumã.

#### Hypostomus chrysostiktos

Bahia State, ANSP 185374, 1, 166.6 mm SL, rio Paraguaçu, rio Paraguaçu basin.

#### Hypostomus commersoni

Uruguay. Montevideo Department. Río de La Plata basin. MNHN A.9444, 425.0 mm SL, holotype, río de la Plata. Brazil. Santa Catarina State, rio Uruguai basin. NUP 15804, 1, 214.0 mm SL, rio Ijuí. NUP 16849, 168.0 mm SL, rio Pelotas. MZUSP 107406, 1, 159.1 mm SL, rio São Francisco, UHE Xingó-CHEESF, downstream the reservoir.

#### Hypostomus delimai

Border of the states of Tocantins and Pará, rio Araguaia basin. NUP 11015, 1, 204.3 mm SL, unnamed stream tributary of rio Araguaia. NUP 11016, 1, 176.7 mm SL, rio Lontra. NUP 11017, 1, 205.5 mm SL, unnamed stream tributary of rio Araguaia.

#### Hypostomus dlouhyi

Paraguay. Alto Paraná Department. Río Paraná basin. MHNG 2229.43, 139.5 mm SL, holotype, río Yguazú.

#### Hypostomusfrancisci

Minas Gerais State rio São Francisco basin. MCP 14038, 1, 180.0 mm SL, Três Marias Reservoir. NUP 9940, 6, 111.0–187.1 mm SL and NUP 9945, 2, 148.6–150.7 mm SL, rio das Velhas.

#### Hypostomus garmani

Minas Gerais State, rio São Francisco basin. BMNH 1904.1.28.3, holotype, 209.9 mm SL; NUP 9819, 9, 87.7–204.2 mm SL; NUP 10028, 1, 78.8 mm SL and NUP 10031, 6, 136.6–170.2 mm SL, all from rio das Velhas.

#### Hypostomus jaguar

Brazil. Bahia State, rio Paraguaçu basin. MZUSP 90870, 13, 68.8–175.6 mm SL, paratypes, rio Paraguaçu, MZUSP 110603, 164.8 mm SL, holotype, rio Paraguaçu. NUP 4448, 2, 126.8–152.9 mm SL, rio Paraguaçu.

#### Hypostomus johnii

Piauí State, rio Parnaíba basin. MCZ 7831, 1, 94.0 mm SL, syntype, rio Poti. MCZ 7864, 2, 93.1–95.5 mm SL, syntypes, rio Poti. NUP 12789, 1, 139.7 mm SL, riacho Quilombo. NUP 12790, 1, 91.2 mm SL, rio Poti.

#### Hypostomus lima

Minas Gerais State, rio São Francisco basin. BMNH 1876.1.10, 2, 72.9–86.1 mm SL, syntypes, Lagoa Santa. NUP5717, 4, 56.1–126.0 mm SL, ribeirão dos Patos. NUP 5721, 2, 47.5–72.8 mm SL, ribeirão das Minhocas. NUP 9827, 18, 81.5–181.5 mm SL, rio São Miguel.

#### Hypostomus macrops

Minas Gerais State, rio São Francisco basin. NUP 9831, 2, 97.7–106.8 mm SL and. NUP 9832, 1, 172.6 mm SL, Rio das Velhas. NUP 9238, 1, 157.9 mm SL, rio Curimataí. *Hypostomus micromaculatus:* Surinam. RMNH 25483,1, 171.0 mm SL, Surinam river. RMNH 25938, 1, 166.0 mm SL.

#### Hypostomus nigrolineatus

rio Jequitinhonha basin, Minas Gerais State: MZUSP 93743, 1, paratype, 115.7 mm SL, rio Araçuaí, municipality of Araçuaí. MZUSP 106743, 2, 192.3–196.5 mm SL, paratypes, municipality of Padre Carvalho, rio Vacaria. NUP 15447, 2, 162.4–212.5 mm SL, paratypes, municipatlity of Grão Mogol, rio Itacambiruçu. NUP 16879, 3, 103.3–138.7 mm SL, paratypes, municipality of Itinga, rio Araçuaí.

#### Hypostomus nudiventris

Ceará State. ANSP 69402, 56.8 mm SL, holotype and NUP 14687, 2, 78.5–100.3 mm SL, rio Choró, municipality of Fortaleza, Northern Brazilian coastal drainages.

#### Hypostomuspantherinus

Bolivia. Beni Departament. AMNH 39946, 2, 128.2–129.5 mm SL, rio Itenez, rio Guaporé basin. Brazil. Mato Grosso State. MCP 35962, 3, 112.8–141.2 mm SL, rio Guaporé, rio Madeira basin.

#### Hypostomuspapariae

Rio Grande do Norte State. ANSP 69398, 94.3 mm SL, holotype, lago Papary, Northern Brazilian coastal drainages. ANSP69399, 1, 99.1 mm SL, paratype, collected with holotype. ANSP 69400, 2, 102.7-126.6 mm SL, paratypes, rio Choró, Northern Brazilian coastal drainages, municipality of Fortaleza. NUP 14684, 10, 54.6–104.4 mm SL, rio Ariri, municipality of Nísia Floresta. *Hypostomuspiratatu:* Paraguay. Paraguarí Department. Río Paraguay basin. MHNG 2265.03, 214.0 mm SL, holotype, río Paraguai.

#### Hypostomusplecostomus

Suriname. MCZ 8025, 1, 169.0 mm SL; exact locality unknown. – RMNH 3102, lectotype (designated by Boeseman, 1968), 221.3 mm SL; Suriname river. – ZMA 105.023, 2, 100.5–110.3 mm SL; Mama creek, Brokopondo.

#### Hypostomuspunctatus

Minas Gerais State. MUP 2605, 2, 172.0–203.0 mm SL, rio Pomba. NUP 9670, 1, 133.3 mm SL, tributary to rio Paraibuna, rio Paraíba do Sul basin. NUP 14483, 1, 220.3 mm SL, rio José Pedro, rio Doce basin. NUP 15488, 5, 117.5-256.6 mm SL, rio José Pedro, rio Doce basin.

#### Hypostomuspusarum

Ceará State, Northern Brazilian coastal drainages. CAS 122225, 142.6 mm SL, holotype, rio Ceará Mirim; CAS 122221, 4, 94.4–141.7 mm SL, paratypes; NUP 14685, 10, 64.7–180.3 mm SL, rio Ceará Mirim, Northern Brazilian coastal drainages. Rio Grande do Norte State, rio Piranhas-Açu basin. NUP 4795, 11, 140.0–207.0 mm SL, rio Acauá and NUP 14683, 2, 103.1–135.0 mm SL, rio Piranhas. Pernambuco State, rio São Francisco basin: NUP 13973, 1, 188.0 mm SL and NUP 13974, 2, 197.5–221.7 mm SL, Itaparica reservoir, rio São Francisco.

#### Hypostomus rhantos

Venezuela. AUM 42100, 4 of 8 paratypes, 161.5–176.8 mm SL; CAS 156859, 1, 70.5 mm SL, río Orinoco. MCZ 68123, 1, 35.0 mm SL. rio Orinoco basin. – LBP 2185, 1, 80.2 mm SL; rio Cataniapo.

#### Hypostomus tapijara

Paraná State, rio Ribeira de Iguape basin. NUP 863, 9, 85.9–251.3 mm SL; NUP 869, 25, 111.0–350.0 mm SL and NUP 2795, 3, 174.9–193.2 mm SL, rio Capivari.

#### Hypostomus unae

Bahia State, rio de Contas basin. NUP 9811, 5, 78.9–53.7 mm SL, rio das Pedras, NUP 9814, 81.5–102.7 mm SL, rio Oricó. MCP 41473, 10, 80.2–126.5 mm SL, rio Preto do Costa. Rio Pardo basin. MCP 41334, 3, 55.2–120.8 mm SL, rio Panelinha.

#### Hypostomus velhochico

rio São Francisco basin. Minas Gerais State: MZUSP 73816, 1, 83.4 mm SL, paratype,municipality of Presidente Juscelino. NUP 12065, 92.6 mm SL, paratype, municipality of Pirapora, rio das Velhas. NUP 12066, 1, 82.8 mm SL, paratype, municipality of Santana do Pirapama, rio das Velhas. NUP 12067, 1, 80.2 mm SL, paratype, municipality of Santana do Pirapama, rio das Velhas.

#### Hypostomus watwata

French Guyana. MNHN A. 8919 (lectotype of *Hypostomus* verres designated by Boeseman, 1968), 194.5 mm SL, Rio Cayenne. Guyana. BMNH 1932.11.10.31 (neotype designated by Boeseman, 1868), 261.2 mm SL, Berbice River.

#### Hypostomus wuchereri

Bahia State. BMNH1863.3.27.15, 1, syntype, 203.8 mm SL, unknown exact locality. BMNH 1852.13.12.8, 1, 127.3 mm SL, syntype, unknown exact locality.

## Acknowledgments

The authors are grateful to Filipe Azevedo (UEM), Rafael Dhovany Marques, João Batista for help in the field collectors. Augusto Frota (UEM) for helping with the map; and Celso Ikedo for helping with fish pictures. Thanks to Barbara Brown and Scott Schaefer (AMNH), Mark Sabaj (ANSP), Patrick Campbell (BMNH), David Catania and Tomio Iwamoto (CAS), Mary Anne Rogers and Kevin Swagel (FMNH), Amanda Cocovicki, Fábio Vieira and Paulo Anchieta Garcia (ICT-UFMG), Lucia Rapp Py-Daniel and Renildo Ribeiro (INPA), Carlos Lucena and Margarete Lucena (MCP), Gilmar Bastos Santos (MCNIP) Claude Weber and Sonia Muller (MHNG), Patrice Pruvost (MNHN), Paulo Buckup, Marcelo Britto and Cristiano Moreira (MNRJ), Ernst Mikschi and Helmut Wellendorf (NMW), Ronald Vonk and Ronald de Ruiter (ZMA) for loan comparative material and hosting museum visits. Nupélia provided logistical support. Capes (Coordenação de Aperfeiçoamento de Pessoal de Nível Superior) provided grants to ISP. We highlight that our experiment was carried out in accordance with the ‘Ethical Principles in Animal Research’ adopted by the Brazilian College of Animal Experimentation (COBEA). The authors declare that they have no conflict of interest.

## Author Contibutions

**Conceptualization:** Cláudio Henrique Zawadzki.

**Data curation:** Cláudio Henrique Zawadzki, Iago de Souza Penido, Tiago Casarim Pessali. Investigation: Cláudio Henrique Zawadzki, Iago de Souza Penido.

**Methodology:** Cláudio Henrique Zawadzki, Iago de Souza Penido, Tiago Casarim Pessali.

**Project administration:** Cláudio Henrique Zawadzki.

**Writing - Original Draft Preparation:** Cláudio Henrique Zawadzki, Iago de Souza Penido, Tiago Casarim Pessali.

**Writing - Review & Editing:** Cláudio Henrique Zawadzki, Iago de Souza Penido, Tiago Casarim Pessali.

